# Multi-Chain Hamiltonian Replica Exchange Framework for Probing Transient Helices and Order-Disorder Transitions in Proteins

**DOI:** 10.64898/2026.10.01.755728

**Authors:** Tika Ram Bhandari, Kurt Kremer, Martin Girard

**Author notes:** Corresponding author’s.

## Abstract

Simulations have emerged as a pillar in biophysics to understand behavior at the molecular scale, in particular for proteins such as intrinsically disordered proteins. These often show transient folding with long-lived states that are challenging to efficiently sample using conventional simulations. Here, we show that Hamiltonian replica-exchange at the level of hydrogen bonds can be used to systematically explore the conformational landscape of proteins. Investigating the IM30 protein, a partially alpha helical protein, under the PLUM coarse-grain force-field, we show that this approach straightforwardly allows characterization of local helix stability, which would be otherwise hard to obtain. For instance, we show that some regions of the conserved coiled-coil structure are less stable than others. By combining the different replicas with principal component analysis, we are able to dissect the conformational landscape of the IM30 protein. Taken together, this indicates that enhanced sampling remains an important part of simulations, allowing insights that cannot be easily obtained in traditional simulations.

**Summary:**

- This work develops a multi-chain Hamiltonian Replica Exchange Molecular Dynamics (HREM) framework using hydrogen-bond strength (HBS) as a tunable parameter to systematically explore protein conformational ensembles and order-disorder transitions.
- A 64-chain dilute solution simulation strategy is introduced to enable efficient and statistically robust sampling of independent protein conformational ensembles within a single simulation box.
- An integrated structural analysis framework is established by combining helicity analysis, polymer physics-based characterization, and principal component analysis (PCA) to quantify secondary-structure formation, transient helix emergence, chain flexibility, and global ensemble organization across different structural states.
- Application of this framework to full-length IM30 and its individual components (H0-3 with helical hairpin and H4-6 IDR) reveals that HBS-dependent order-disorder transitions in all constructs are governed by transient helix formation. In the full-length protein, IDR-associated transient helices reshape the conformational landscape of the conserved hairpin architecture, enabling a continuum of conformational states through dynamic ensemble reorganization.
- This work demonstrates that IDRs and their associated transient helices act as key regulators of protein conformational organization, promoting continuous transitions between ordered and disordered states and providing a general framework for investigating conformational dynamics in modular proteins.

## Introduction

The helix is a ubiquitous structural motif in proteins and nucleic acids, representing a precise stereostructural arrangement that imparts distinctive physical, chemical, and mechanical properties [1–5]. Helical macromolecules often behave as rigid-rod polymers with long persistence lengths [6, 7] and can self-assemble into diverse nanostructures, including fibers, vesicles, networks, and helical phases [8]. In proteins, structured helical segments frequently coexist with intrinsically disordered regions (IDRs), flexible segments that lack stable secondary structure but play essential roles in protein function [9–12]. This combination gives rise to modular architectures in which ordered helical segments are interspersed with disordered regions, forming organizations such as IDR-helix-IDR or helix-IDR-helix, as illustrated in Figure 1A.

**Figure 1:**
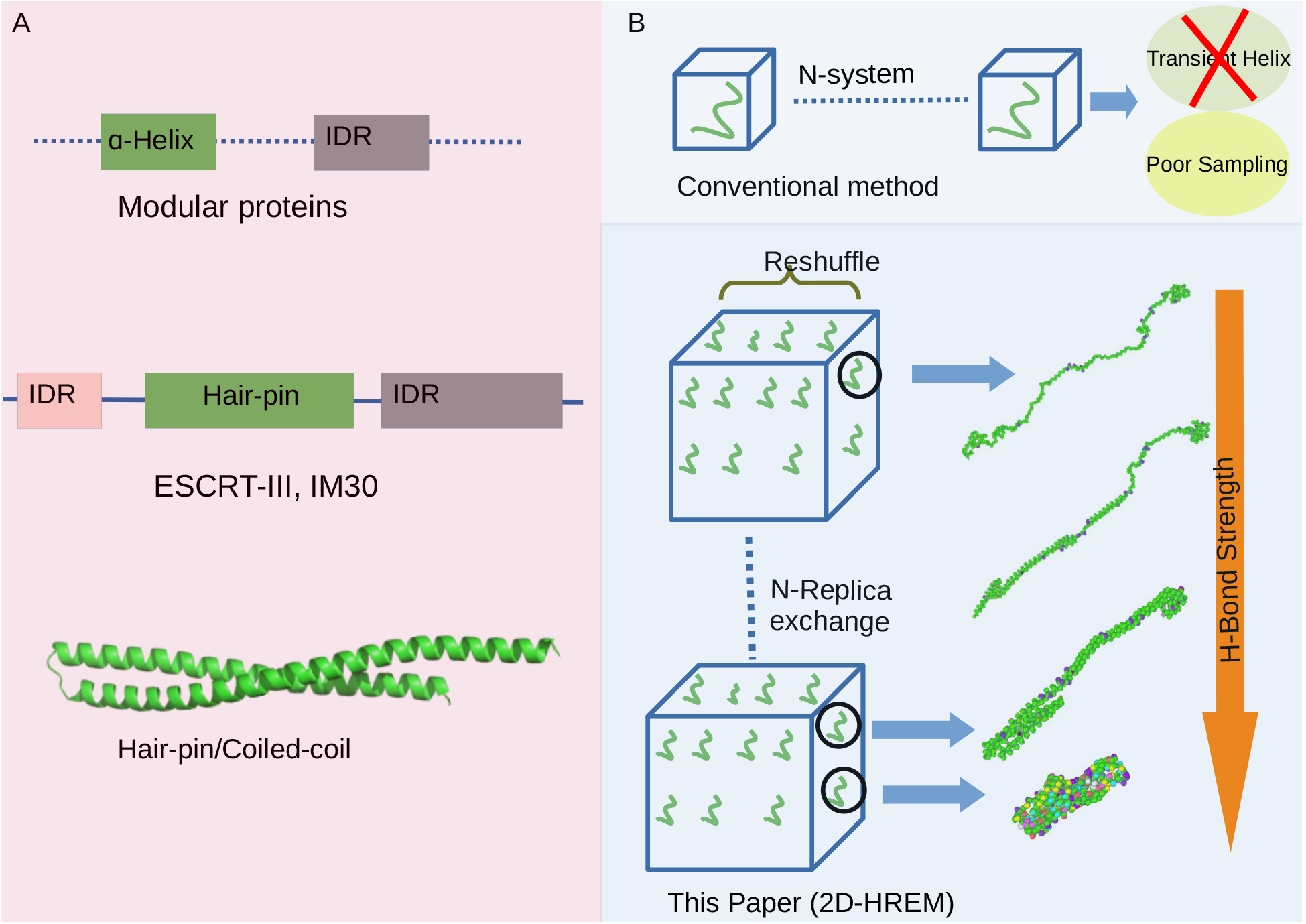
(A) Schematic representation of the modular architecture of ESCRT-III, IM30 protein, highlighting the conserved helical hairpin core flanked by intrinsically disordered regions (IDRs).(B) Schematic illustration of the Hamiltonian Replica Exchange Molecular Dynamics (HREM) framework used in this study. The upper panel depicts the conventional simulation strategy, in which a single protein chain is simulated per simulation box without replica exchange, resulting in limited conformational sampling and poor characterization of transient helical states. The lower panel illustrates the proposed HREM approach, where multiple noninteracting protein chains are simulated simultaneously in a dilute environment while systematically varying the HBS across Hamiltonian replicas. This strategy enables efficient exploration of conformational space, enhanced sampling of folding-unfolding transitions, and reliable characterization of transient helical conformations that are difficult to capture using conventional simulations.

Proteins typically consist of compact, independently folded domains connected by flexible linkers, while helical motifs composed of two to four helices can form hairpin or coiled-coil structures that provide additional structural stability [13, 14]. In contrast, IDRs are generally enriched in polar and charged residues, such as Gln, Ser, Pro, Glu, and Lys, and depleted in bulky hydrophobic residues, enabling dynamic interactions and functional versatility [15, 16]. Thus, proteins can occupy a broad continuum of structural states, ranging from well-folded globular domains to molten globules, partially disordered or highly flexible regions, and fully unstructured chains [11, 12, 32–34].

A wide range of experimental techniques has been developed to probe protein structure and dynamics, including X-ray crystallography, circular dichroism, Raman and fluorescence spectroscopy, hydrodynamic measurements, proteolytic assays, and small-angle X-ray scattering (SAXS) [17–24]. While crystallography can resolve structures at high spatial resolution, it is not well suited to highly flexible or IDRs [12]. SAXS, in contrast, provides low-resolution structural information in solution and is particularly valuable for characterizing conformational ensembles, domain motions, and intrinsically disordered proteins (IDPs) [23, 24]. Nuclear magnetic resonance (NMR) spectroscopy and molecular simulations further complement these approaches by providing information on the dynamic ensembles and folding pathways of IDPs [25–31].

Despite lacking stable hydrophobic cores, IDPs can adopt well-defined tertiary structures upon binding to interaction partners through a process known as coupled folding and binding. During this process, transient secondary-structure elements, including helices, may form but often remain only marginally stable and can be readily perturbed by changes in sequence or environment [35]. In multidomain proteins, IDRs introduce additional conformational heterogeneity that can enhance the structural diversity of neighboring conserved domains. Although IDPs and IDRs have been extensively studied, accurately characterizing their heterogeneous conformational ensembles and transitions between ordered and disordered states remains a major computational challenge. In particular, conventional molecular dynamics simulations can face difficulties in efficiently sampling transient secondary structures and the broad conformational landscapes associated with proteins containing both structured and disordered regions. Computational approaches that enable controlled exploration of these ensembles are therefore essential for understanding how intrinsic disorder influences protein structure and dynamics.

During folding-unfolding transitions, protein segments do not necessarily undergo complete unfolding. Instead, they can transiently form helices or helical segments of varying lengths that continuously interconvert between folded and unfolded states, reflecting their dynamic and short-lived nature [36]. These transient structures can modulate local order-disorder transitions and influence the overall conformational landscape of a protein [37–39]. Previous studies have extensively investigated helix-coil transitions [40, 41] and protein folding pathways [42]. However, the physical origins, functional roles, and dynamical consequences of transient helices in heterogeneous protein sequences remain poorly understood.

Much of the existing work has relied on simplified homopolypeptide models, which tend to exhibit relatively uniform folding and unfolding behavior and therefore have limited ability to reproduce the heterogeneous helicity observed in natural proteins. Related studies have also examined simple model systems, including elastin-like peptides [43–45]. By comparison, transient helices in IDPs remain less well characterized. A notable example is the transient helix in the TDP-43 low-complexity domain (LCD) [46–48], which has been implicated in phase separation, functional interactions, and the formation of dynamic, partially ordered structural ensembles. Capturing such transient and heterogeneous helicity requires model sequences that combine ordered and disordered regions. Experimentally, sequences capable of transient helix formation can remain predominantly disordered under physiological conditions. Designing and characterizing such sequences is therefore challenging because they must simultaneously balance local structural stability and conformational flexibility [49, 50].

Molecular simulations can provide detailed insight into these transitions, but atomistic simulations of large or heterogeneous systems are computationally demanding [51]. Sequences containing multiple structured regions may require long simulations and multiple replicas to achieve adequate equilibration and statistically meaningful sampling [52, 53]. Consequently, many computational studies of helix-coil transitions have employed simplified systems such as polyalanine, often using relatively short atomistic simulations to investigate helix formation and unfolding [40, 41]. Although these approaches provide valuable information about intrinsic helix stability, their limited sequence complexity can restrict their ability to capture the full spectrum of folding-unfolding configurations present in heterogeneous proteins.

Helix-coil transitions can be induced by changes in temperature or pressure that perturb secondary-structure stability [54, 55]. However, such approaches primarily perturb the thermodynamic stability of hydrogen-bonded secondary structure and may not provide direct control over the local interactions governing helix formation. A suitable model for studying folding-unfolding transitions should therefore incorporate heterogeneous sequences capable of supporting modular secondary structure while remaining computationally tractable. Coarse-grained (CG) models provide an effective framework for this purpose because they enable efficient sampling of broad conformational landscapes over extended timescales and for larger systems [51, 56–59].

Here, we employ the PLUM force field, a physics-based and transferable CG model that represents amino acids using four beads per residue (three for glycine) within an implicit-solvent framework and includes an explicit hydrogen-bond potential. Unlike Go-like models [60, 61], PLUM does not require prior knowledge of the native structure. Instead, it incorporates both local interactions and non-native contacts, allowing *α*-helices and *β*-sheets to form, fluctuate, and rearrange spontaneously. This balanced representation of secondary structure enables efficient sampling of conformational transitions and long-timescale protein dynamics [62, 63]. By systematically varying the hydrogen-bond strength (HBS) while integrating concepts from polymer physics and principal component analysis (PCA), we investigate protein folding-unfolding transitions with particular emphasis on the emergence of transient helicity and its role in shaping protein dynamics and conformational ensembles.

In this study, we investigate IM30, a member of the ESCRT-III superfamily involved in membrane remodeling and the assembly of higher-order protein complexes [64]. Members of the ESCRT-III family share a conserved structural architecture consisting of a central helical hairpin domain flanked by IDRs, which provide structural flexibility and functional adaptability. In bacterial homologs such as IM30, an additional IDR is present at the N terminus, whereas eukaryotic ESCRT-III proteins typically contain only a C-terminal IDR, as illustrated in Fig. 1A. The structural organization of the IM30 monomer, extracted from the barrel structure (PDB: 7O3Y), is shown in Fig. 2A. IM30 comprises 267 residues (Fig. S9) arranged into seven helical segments (H0-H6) connected by flexible linkers. In the assembled barrel, these helices are well defined, whereas in the disassembled state several helices partially unfold or disappear, reflecting IM30’s intrinsic tendency to reside near an order-disorder transition [17]. The protein contains a rigid coiled-coil/hairpin core (H1-3) flanked by a short N-terminal helix (H0) and a highly dynamic C-terminal region (H4-6) [17, 64]. The residue boundaries of the individual helical segments are summarized in Fig. 2B.

**Figure 2:**
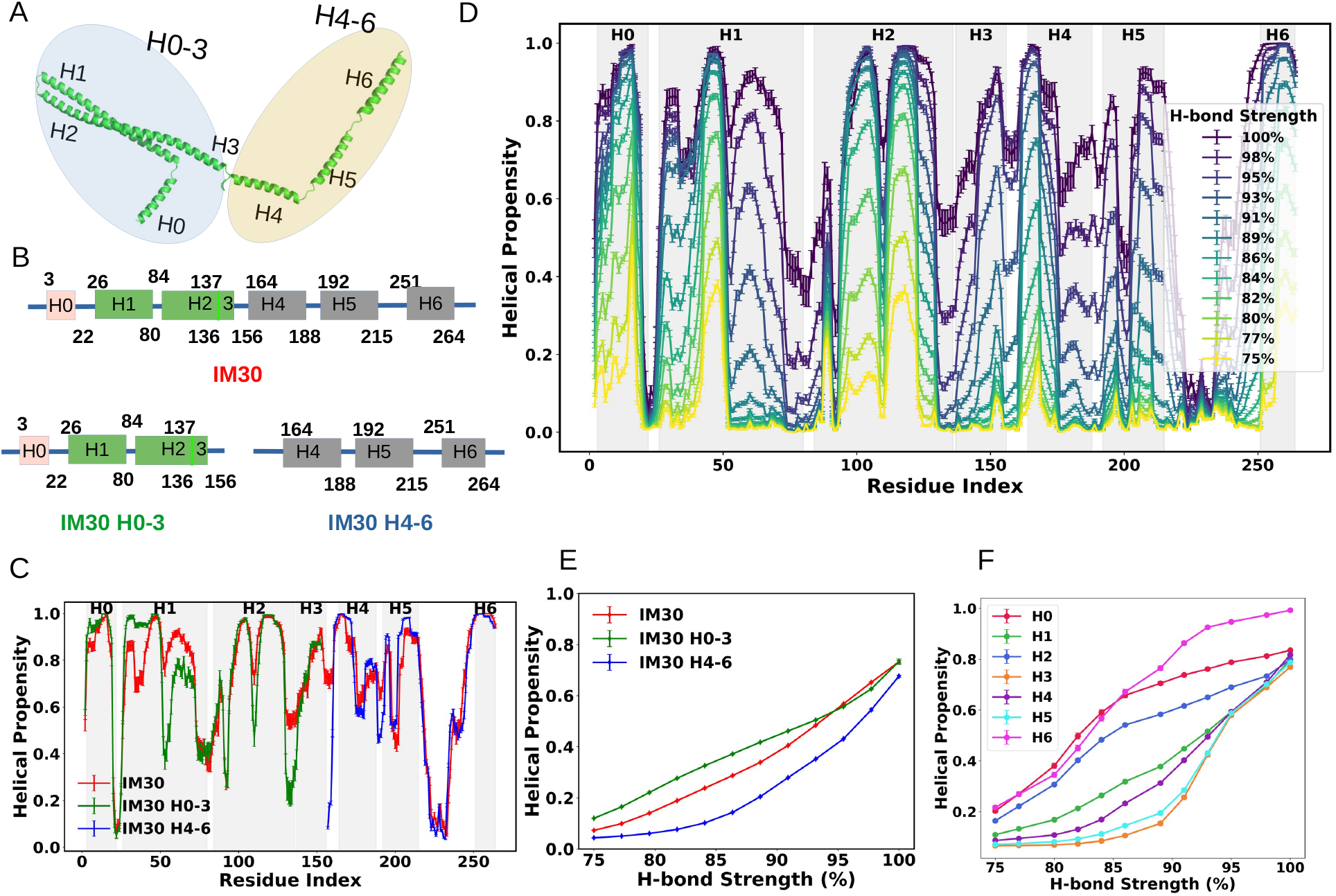
(A) Structure of IM30 showing its seven helical segments (H0-H6). Segments H0-3 form the coiled-coil region, while H4-6 constitute the C-terminal region. The structure represents a monomeric unit of the IM30 barrel (PDB: 7O3Y).(B) Schematic block representation of the helical segments with corresponding residue ranges. (C) Helical propensity of IM30, H0-3, and H4-6 at the residue level at 100% HBS, shown in red, green, and blue, respectively. (D) Residue-wise helical propensity of IM30 across varying hydrogen-bond biases. Darker colors indicate higher HBS (100%), while lighter yellow indicates lower strength (75%). (E) Comparison of the average helical propensity of IM30, H0-3, and H4-6 across different HBSs. (F) Helical propensity profiles for individual helical segments H0-H6 of IM30. *Error bars in all plots represent standard deviations*.

The interplay between the ordered hairpin and the disordered terminal regions is central to IM30 function, regulating its ability to assemble into higher-order oligomers and biomolecular condensates under similar environmental conditions [65]. While the coiled-coil hairpin remains relatively stable, the flanking regions undergo substantial conformational fluctuations [17]. To retain the native structural context while enabling efficient exploration of the conformational landscape, we consider three complementary constructs (Fig. 2B): (i) the H0-3 construct, containing the N-terminal region and coiled-coil hairpin (156 residues); (ii) the C-terminal H4-6 construct (111 residues); and (iii) the full-length IM30 monomer (267 residues). Comparison of these three systems allows us to distinguish the intrinsic conformational behavior of the structured and disordered regions from their coupling within the intact protein.

To explore their conformational ensembles, we employ CG Hamiltonian Replica Exchange Molecular Dynamics (HREM), in which HBS is systematically varied while all other interaction parameters remain unchanged. Modulating HBS provides a physically motivated means of tuning secondary-structure stability and efficiently sampling the order-disorder landscape. By comparing the H0-3, H4-6, and full-length constructs across a range of HBS values, we quantify how local hydrogen-bond interactions regulate helicity and how these local structural changes propagate into global conformational transitions through transient helices.

## Results

We first establish a simulation framework that enables efficient sampling of multiple independent protein molecules while preserving dilute-solution conditions. We then show that systematically tuning the HBS not only modulates protein rigidity by controlling helical stability, but also provides direct access to transient helix formation and continuous order-disorder transitions. Exploring these transitions is essential because they determine the conformational ensemble accessible to the protein and thereby its capacity for structural and functional organization. By comparing the isolated H0-3, the H4-6, and the full-length IM30 monomer, we directly assess how the intrinsically disordered flanking regions reshape the conformational landscape of the conserved ESCRT-III hairpin. This comparison provides a mechanistic framework for understanding how a structurally conserved core can acquire rearrangement through flexible regions.

Here, the PLUM coarse-grained (CG) model represents hydrogen bonding implicitly through a 12-10 Lennard-Jones potential combined with an orientation-dependent angular term (Eq. 1) [62, 66]. To systematically tune secondary-structure stability, the hydrogen-bond interaction strength was scaled from 75% to 100% of its reference value using 12 Hamiltonians parameterized by the scaling factor, *λ*. This controlled modulation of hydrogen-bond interactions provides a computational approach to perturb secondary-structure stability and explore conformational transitions analogous to those observed under denaturing conditions, such as urea-induced destabilization [17]. Enhanced sampling across this Hamiltonian space was achieved using HREM [67]. For each of the three IM30 constructs, 64 protein chains were simulated simultaneously within a single dilute simulation box. The N-terminal bead of each chain was restrained to a fixed anchor position, ensuring that individual molecules remained spatially separated throughout the simulation and thereby eliminating inter-chain interactions by construction. Consequently, each chain sampled its conformational ensemble independently, allowing a single simulation to generate 64 statistically independent trajectories while retaining the computational efficiency of a multi-chain framework. Each trajectory was propagated for 20 *µ*s.

A schematic overview of the simulation strategy is presented in Figure 1B. The figure illustrates the multi-singlechain simulation setup, representative protein conformations spanning low to high HBS, and the two-dimensional HREM workflow. Compared with conventional approaches that require numerous independent simulation boxes, our strategy provides substantially improved sampling efficiency by simultaneously exploring conformational and Hamiltonian space within a single framework. This enhanced sampling is particularly advantageous for capturing rare, transiently populated helical states that are often inaccessible in conventional molecular dynamics simulations. Complete details of the simulation protocol are provided in the Methods section and Supporting Information.

### Helical Spectra Reveal the Formation of Transient Helices

Helical propensity analysis provides a measure of the protein’s degree of disorder. The backbone dihedral angles (*ϕ, ψ*) within the *α*-region of the Ramachandran plot (Figure S4 A) were used to define residue helicity. A residue is classified as helical if its dihedral angles satisfy −160° *≤ ϕ ≤ −* 20° and −120° *≤ ψ ≤* 50°. A residue is considered part of an *α*-helix when it is in a helical conformation, and both its neighboring residues are also helical. These criteria are simulation-based and adapted from previously established definitions of helical states, ensuring consistency with prior computational studies. The helical propensity of a residue is then defined as the fraction of simulation time during which it forms part of an *α*-helix [17, 68, 69]. The residue-level helical propensity of IM30 are shown in Figures 2C and 2D. The helicity of IM30 (red), H0-3 (green) and H4-6 (blue) at 100% HBS in Figure 2C illustrate coiled-coil region of IM30 is highly structured and it is separated by H0 through a kinks represented by sharp decrease of propensity for the residues around 20-25 also matched with PDB structure. Further, with in coiled-coil, the IM30 also has distinct disorder region around residue 80-90, which connects H1 and H2. Similarly, regions which corresponds other connectors shown by 2A and 2B are also found in the helicity profile for IM30. The helicity of H0-3 and H4-6 from independent simulation is compared with IM30. The results for both are mostly overlapping with IM30, except that the region corresponds to H1 of the coiled-coil and N-terminal of H4-6. The drop of helicity of N-terminal in H4-6 indicates that some residues (such as QAA at C-terminal of H3) on IM30 act as agents to stabilize the helicity of H4-6 in full length.

Helicity profiles of IM30 for all biased HBS values are shown in Figure 2D. Dark colors indicate 100% helicity (matching the red profile in Figure 2C), while lighter colors represent lower helicity. The shaded areas in the plots correspond to these segments, highlighting residues most prone to adopting *α*-helical conformations under different H-bond biases. Residues in the linkers between helical segments generally exhibit lower helical propensity than those in structured regions. For example, the kinks between H0 and H1 and the connector between H5 and H6 remain largely unstructured across all HBS values. Within the helical segments, however, stability is heterogeneous: some regions retain helicity while others unfold more readily as HBS is reduced, highlighting that even largely helical regions can exhibit local differences in folding propensity. Structured regions can form helices even at low HBS and are more responsive to increases in HBS, folding more readily than linkers or other disordered regions. Figure 2D shows that H0, H1, H2, and H6 contain residues with high propensity to form ordered structures, mostly in the middle of each segment, whereas H3, H4, and H5 form helices only at high HBS. These three segments are mostly disordered following barrel disassembly of IM30. Capturing such differing folding propensities of residues and regions using our method enables the study of order-disorder transitions in modular proteins such as IM30. Similar patterns are observed for H0-3 and H4-6 in their corresponding regions of IM30, with the exception of H1: in the isolated fragment, the first half of H1 has higher helicity, whereas in the full-length protein, the second half is more helical. This variation along the helix suggests that different subregions may have distinct folding properties, depending on residue contacts within the tertiary structure (snapshots in Figure S2A) in the full-length protein compared to isolated fragments. However, the overall folding capacity of residues in these independent fragments is lower, as seen in Figure 2E. The helicity profiles for H0-3 and H4-6 under different HBS values are shown in Figures S4B and S4C, respectively. To gain insight into fragment helicity in full-length IM30 and its isolated segments H0-3 and H4-6, we calculated the average helicity of each segment. Each segment exhibits distinct folding ability, as shown in Figures 2F, S4E, and S4F. Helicity profiles and slopes indicate that H6 generally has the highest propensity and can form a helix even at low HBS, whereas H3, H4, and H5 are less likely.

The distinct folding behaviors of H1 arise from sequence heterogeneity and segment length. Helix-propensity values (according to the Pace and Scholtz scale [70]) indicate that Alanine (A) has the minimum value (0), representing maximal *α*-helix propensity, whereas Glycine (G) has the maximum value (1), representing minimal helix propensity and increased flexibility. When the sequence of H1 (highlighted within IM30 in Figure S9) is divided into two halves (residues 1-30 and 31-60), they exhibit distinct characteristics. The first half (EKVLEQAVIDMQEDLVQLRQAVARTIAEE) contains several clusters of residues with low helix-propensity values, such as LEQ, QAV, and EQAV, enriched in hydrophobic and moderately charged residues (L, V, A, Q, E). These clusters are likely to stabilize *α*-helical structures and contribute to the rigidity of this segment. In contrast, the second half (KRTEQRLNQDTQEAKKWEDRAKLALT) is enriched in polar and charged residues (N, D, T, E, K) and contains fewer residues with low helix-propensity values. Triplets such as NQD and DTQ have relatively high helix-propensity values on this scale, indicating that they are less likely to form *α*-helices and are therefore more flexible or disordered. This analysis illustrates that sequence context can strongly influence folding. This behavior is consistent with the well-established principle that local sequence composition and context influence secondary structure propensity and protein conformational preferences [12, 70, 71]. Consequently, regions with high intrinsic disorder propensity may adopt structure within the full-length protein due to stabilizing tertiary contacts, even if they remain unfolded in isolation. Conversely, highly ordered segments that fold in isolated fragments may become partially destabilized or unfold in the context of the full-length protein, for example, due to competing intramolecular contacts or local environmental effects. Such behavior could arise from cooperative folding-unfolding upon binding [72, 73], which is beyond the scope of the present study. Depending on the sequence, transitions may involve *α*-helix → *β* → coil conversions [74].Thus, the folding propensity of a segment varies with its chemical composition, which influences the formation of intramolecular tertiary contacts. In addition to this general behavior, IM30 exhibits pronounced region-dependent secondary structure characteristics, including transient *α*-helical formation in specific segments. These transient helices display varying stability and are sensitive to local sequence context as well as intramolecular interactions, which modulate their persistence and degree of helicity. Overall, this sequence-dependent stabilization leads to heterogeneous structural behavior along the chain and contributes to the conformational variability of IM30. These observations are consistent with previous studies showing that transient helices in disordered or modular proteins can be stabilized through local sequence effects and intramolecular interactions [71, 75, 76].

Furthermore, the average helicity graphs of the four H0-H3 segments and the three H4-H6 segments extracted from full-length IM30 (Figure S4D) reveal a common transition point at approximately 93% HBS for IM30, H0-H3, and H4-H6. This value corresponds to the recommended HBS for disordered regions in the PLUM force field, as it balances secondary structure formation with realistic sampling of disorder [77]. Below this threshold, residues in structured regions dominate helicity, while above 93% HBS, disordered regions begin to adopt helical conformations, creating additional strain on residues with high folding propensity. Consequently, the summed helicity of H0-H3 decreases relative to H4-H6, reflecting segmental interactions within tertiary structure. Beyond this point, the protein exhibits increased helicity, as the PLUM force field promotes secondary structure formation, driving self-organization into tertiary arrangements resembling those observed in PDB structures.

### Modular Proteins Exhibit Semi-Flexible Polymer Behavior

Helical spectra analysis presented in the previous section revealed heterogeneous residue-level structural stability within the protein sequence, where different residues display distinct propensities for stable helices, transient helices, or weakly populated helical states. Such variation in local secondary-structure stability reflects the modular organization of proteins, where structured and disordered regions coexist. To characterize the resulting global chain behavior, polymer physics provides a valuable framework for describing the conformational properties of protein chains. In the long-chain limit of homopolymers, polymer physics predicts scaling relations such as *R*(*s*) *∼s*^*ν*^, where *s* is the contour separation between two monomers (or residues in our case), and the exponent *ν* characterizes chain behavior. Specifically, *ν* = 0.5 corresponds to ideal chains in a *θ*-solvent, *ν ≈* 0.59 to swollen chains in good solvent conditions, and *ν ≈*1*/*3 to collapsed chains in poor solvent environments [78]. These relations strictly apply in the asymptotic long-chain limit. For finite-sized proteins, however, deviations from ideal scaling are expected due to limited chain length, structural constraints, and sequence heterogeneity. Unlike homopolymers, proteins are heteropolymers whose conformational behavior is influenced by sequence-specific interactions. Variations in hydrophobicity and residue composition modulate effective intrachain interactions and can be interpreted as altering the effective solvent quality in a sequence-dependent manner. Thus, while the scaling framework remains useful, the effective exponent *ν* reflects both environmental conditions and intrinsic sequence properties. The interpretation of *R*(*s*) scaling must be placed in the context of flexible versus semi-flexible polymer behavior. For a flexible polymer in a collapsed globular state, one expects *R*(*s*) *∼ s*^1*/*2^ at short contour separations, followed by a plateau corresponding to the finite size of the globule [79–81]. In contrast, a semi-flexible polymer exhibits rod-like scaling (*R*(*s*) *∼ s*) for *s < l*_*p*_, where *l*_*p*_ is the persistence length, and crosses over to flexible behavior for *s > l*_*p*_.

The log-log plot of the internal distance *R*(*s*) as a function of segmental separation *s* for IM30, H0-3, and H4-6 at 100% HBS is shown in Figure 3A (red, green, and blue curves, respectively). Here, *R*(*s*) denotes the average spatial distance between two residues separated by *s* positions along the sequence. Results for other bias values are provided in Figure S5. At low HBS, the observed scaling behavior for all systems is consistent with a compact globular polymer: the slope approaches approximately 1*/*2 at intermediate *s*, followed by a plateau, indicating finite-size collapse. This behavior is compatible with standard polymer expectations. At higher HBS, the slope at small *s* increases, reflecting enhanced local stiffness due to helix formation. This corresponds to an increase in the effective *l*_*p*_. However, the large-*s* regime continues to exhibit plateau behavior, indicating that the overall conformation remains globule-like. Thus, increasing HBS primarily modulates local stiffness rather than inducing a qualitative change in global polymer architecture.The crossover near *s ≈ l*_*p*_ marks the transition from rod-like to flexible behavior. The regime *s < l*_*p*_ was fitted linearly to quantify local stiffness (Figure 3A, inset (a)). The resulting slopes indicate the order of rigidity as IM30 (0.76) *>* H0-3 (0.73) *>* H4-6 (0.70) for 100% HBS. All slope values remain below unity, consistent with semi-flexible polymer behavior rather than rigid-rod scaling. Here, the full-length IM30 construct exhibits greater rigidity than the isolated helical domains despite displaying similar overall helical propensities. This suggests that multivalent interactions within the full-length protein architecture enhance helix stabilization and contribute to increased structural rigidity. The *l*_*p*_, expressed in units of number of residues and independently estimated from angular correlation analysis (Figure 3B), follows the same trend, with values of 8.60 for IM30, 6.82 for H0-3, and 6.13 for H4-6 under 100% HBS conditions. Since helices introduce locally rigid segments, *l*_*p*_ should be interpreted as an effective, sequence-averaged measure rather than a uniform mechanical property along the chain [82].

**Figure 3:**
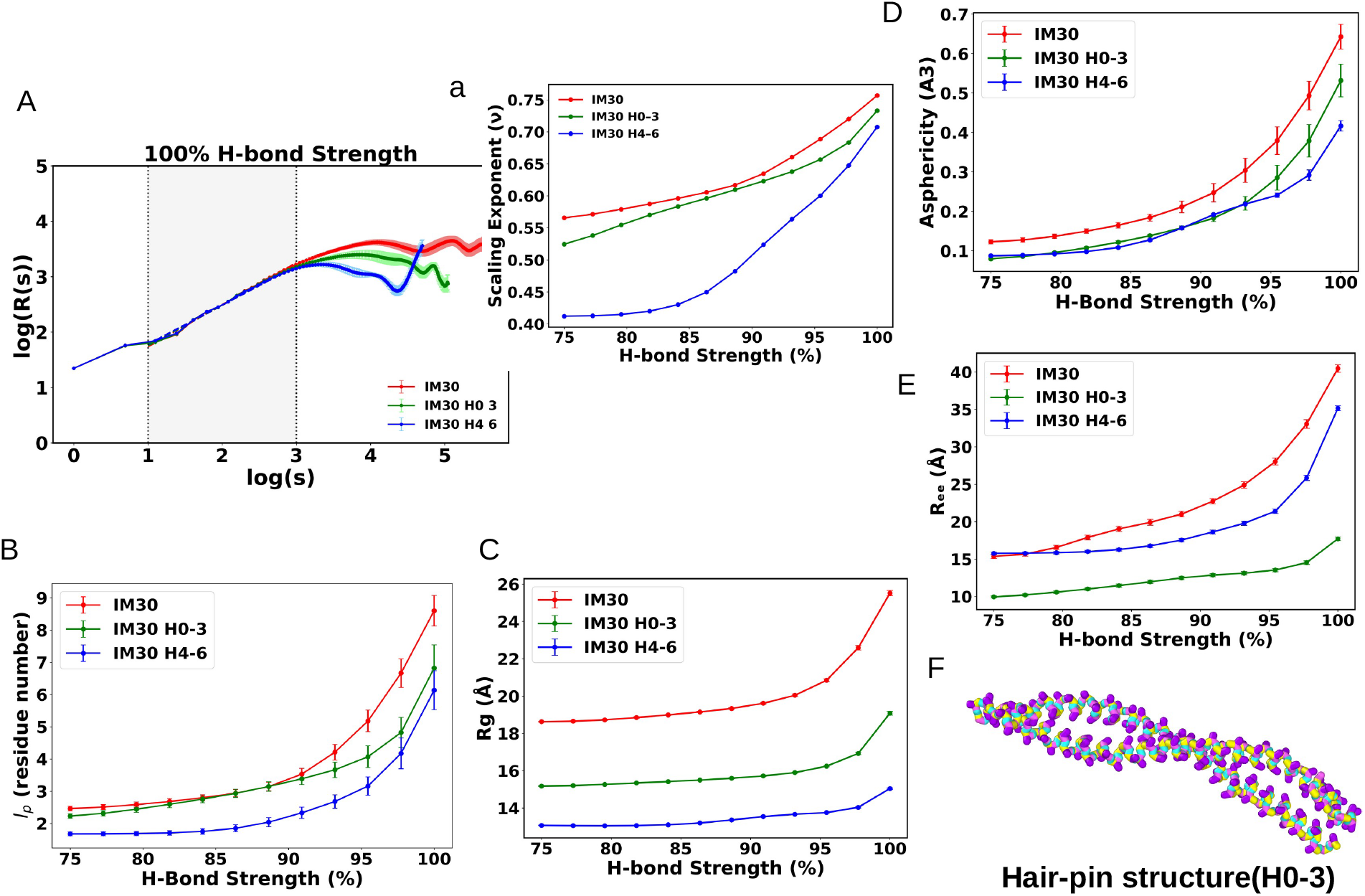
(A) Scaling relation of *R*(*s*) with residue spacing for HBS of 100% and inset shows the slope of the scaling curves for all chains and all bias strengths, obtained from a linear fit over the range 1 *≤* log(s) *≤* 3. (B) shows the *l*_*p*_ calculated from the angular correlation for IM30, H0-3, and H4-6 (C-E) *R*_*g*_, *A*3, and *R*_*ee*_ for IM30, H0-3, and H4-6. Red, green, and blue represent full-length IM30, H0-3, and H4-6, respectively. (F) Snapshot of H0-3 illustrates hair-pin like structure.*Error bars in all plots represent standard deviations*.

To further characterize global conformational properties, we computed the radius of gyration, *R*_*g*_, the asphericity, *A*_3_, and the end-to-end distance, *R*_*ee*_. The corresponding results are shown in Figures 3C-E. The *R*_*g*_ values follow the order IM30 *>* H0-3 *>* H4-6 across all HBS strengths, reflecting differences in chain length and overall mass distribution. The *A*_3_ provides insight into global shape anisotropy. IM30 consistently exhibits larger *A*_3_ values than the fragments, indicating a more elongated overall shape. H0-3 and H4-6 display comparable *A*_3_ values below 93% HBS, suggesting similar global anisotropy in this regime. The *R*_*ee*_ (Figure 3E) is smallest for H0-3 across all biases, indicating stronger long-range compaction relative to IM30 and H4-6. Taken together with the *R*(*s*) scaling, *l*_*p*_, and shape descriptors, these observations suggest that H0-3 adopts a highly organized intrachain arrangement, with residues folding back onto each other in a hairpin-like manner. To illustrate this, a representative simulation snapshot showing the hairpin configuration is provided as Figure 3F, allowing direct visualization of the corresponding intrachain contacts.

Interestingly, *l*_*p*_ of IM30 and H0-3 coincides for HBS values below approximately 93%, suggesting that both systems exhibit comparable effective stiffness in this regime. This HBS is consistent with previous studies on IDPs [77], which showed that the H-bond parameter must be scaled to approximately this value to reproduce experimental observations. This observation further indicates that at 93% HBS, only the hairpin of IM30 remains ordered and contributes to plasticity, whereas helices H0 and H4-6 likely continue to unfold or their contribution is insignificant. This behavior is consistent with experimental observations showing that, following barrel disassembly, only the central hairpin remains structured in monomeric IM30 [17, 83]. Above 93% HBS, however, the *l*_*p*_ curves begin to diverge, with IM30 exhibiting a more pronounced increase. This trend suggests that the IDRs progressively undergo folding at higher HBS, locally rigidifying the backbone and thereby increasing the effective stiffness of the full-length protein. Such an increase in rigidity is also reflected in the helical propensity (Figure 2E): for IM30, the slope becomes steeper above 93% HBS, whereas the H0-3 construct displays a more uniform trend across the HBS range. In contrast, the *A*_3_ shows that H0-3 and H4-6 coincide in the low HBS regime, indicating comparable global shape anisotropy under weak HBS. This similarity likely reflects, at least in part, their comparable chain lengths (156 and 111 residues, respectively), which constrain the accessible conformational space in a similar manner at low stiffness.Taken together, these observations suggest that below *∼* 93% HBS, H0-3 behaves predominantly as a flexible collapsed polymer. Above this value, however, it increasingly deviates toward semi-flexible behavior, as reflected by the separation in *l*_*p*_ and scaling characteristics. Notably, a HBS of *∼* 93% also emerged in the helicity analysis as the regime where secondary structure becomes significantly stabilized in H4-6, a predominantly disordered protein region. This value defines a crossover region separating a more flexible, weakly ordered regime from a stiffer, more locally ordered state.

### Structural Stability of the Conserved Helical Hairpin

All ESCRT-III family members contain a highly conserved helical hairpin motif (Figure 1A) [64]. This hairpin architecture has been shown to exhibit remarkable stability, as demonstrated during the barrel disassembly of IM30. Moreover, the IM30* variant retains the same conserved hairpin structure, which remains stable up to a defined range of urea concentrations, as reported in our previous work [17]. The same study also demonstrated that the isolated H0-3 fragment can independently adopt a hairpin-like structure. Furthermore, recent work has shown that this conserved motif remains stable during membrane association and in membrane-carpet-like assemblies [20]. Consistent with these experimental observations, our simulations reveal that the conserved hairpin motif adopts distinct structural conformations, as characterized through polymer physics-based structural analysis. Although the simulations were initiated from disordered conformations using a physics-based model, several sampled structures (snapshots shown in Figure 3F) exhibit conformations consistent with experimentally observed hairpin architectures (Figure 1A). In addition, the overall end to end distance (*R*_*ee*_) distributions (Figure 3E) indicate the presence of compact, supertertiary-like conformational states associated with this conserved domain. Interestingly, comparison of the full-length protein with isolated hairpin-containing fragments and C-terminal fragments reveals that these conserved structural elements undergo conformational rearrangements within the intact protein architecture. This changes likely arises from the influence of flexible flanking regions, which modulate the conformational landscape of the conserved motif while preserving its overall structural integrity. Therefore, exploring the complete conformational ensemble is essential for understanding how conserved structural elements adapt within modular proteins such as ESCRT-III, where ordered domains coexist with IDRs. The role of these flexible regions in regulating hairpin conformation is discussed in the following section.

### Transient Helices Shape a Conformational Landscape

In the previous section, we analyzed the helicity and structural properties of IM30 and its fragments, showing that tertiary chain compaction and semi-flexible behavior persist even at highest HBS value and correlate with the formation of transient helices. Regions with high intrinsic helical propensity contribute to local structural ordering, whereas more disordered segments behave as flexible polymers and acquire helical structure only at elevated HBS. These observations reveal how spatially heterogeneous helicity drives an order-disorder transition: local helices transiently stabilize individual segments without imposing a rigid global fold. The crossover near *∼* 93% HBS, observed consistently from both helicity and polymer analyses, represents a regime where disordered regions begin to acquire significant local order, highlighting the dynamic coupling between transient secondary structure formation and global chain compaction.

To investigate how these structural changes are reflected in the conformational ensembles, we performed principal component analysis (PCA) of IM30, H0-3, and H4-6. The PCA was constructed using a combined feature space consisting of segmental helicity and inter-segment center-of-mass (CoM) distances (seven helicity features and 21 distance features for IM30), thereby capturing both local secondary-structure formation and global conformational organization. Equivalent feature sets were generated for the isolated fragments. For each system, the resulting ensemble contained 64 chains × 10,000 frames × 12 HBS conditions, providing extensive sampling of the conformational landscape. In all systems, the principal components capture coupled variations in helicity and inter-segment distances. PC1 is primarily associated with changes in helicity and structural expansion, whereas PC2 reflects the anticorrelation between inter-segment distances and helicity, with strong contributions from CoM distance variations (Figure S7). Notably, the IDR-associated distances (D4-6 and D5-6) contribute negatively to PC1, reflecting the distinct behavior of flexible regions compared with structured helical segments. Together, these PCs effectively describe the coupled changes in local ordering and global architecture. The broad distribution with low variance % in PCA space reflects the extensive conformational heterogeneity sampled by the simulations, consistent with the dynamic nature of IM30 and its disordered regions.

To provide a structural overview of the helical conformational states, the left panels of Figure 4A-C show the AlphaFold-predicted structures of the corresponding protein constructs, whereas the right panels present the PCA projections of their simulated conformational ensembles. The predicted structures of the isolated domains retain similar helical characteristics to the full-length protein, suggesting that these regions preserve their intrinsic structural propensity even in the absence of the complete architecture. Notably, the predicted structures contain a flexible region between helices H5 and H6, consistent with the reduced helical propensity observed in our simulation-derived helicity profiles (Figure 2D). IM30 and H0-3 exhibit pronounced shifts in PCA space with increasing HBS, progressing toward regions characterized by increased inter-segment distances and enhanced helicity, corresponding to folded-expanded conformational states.

**Figure 4:**
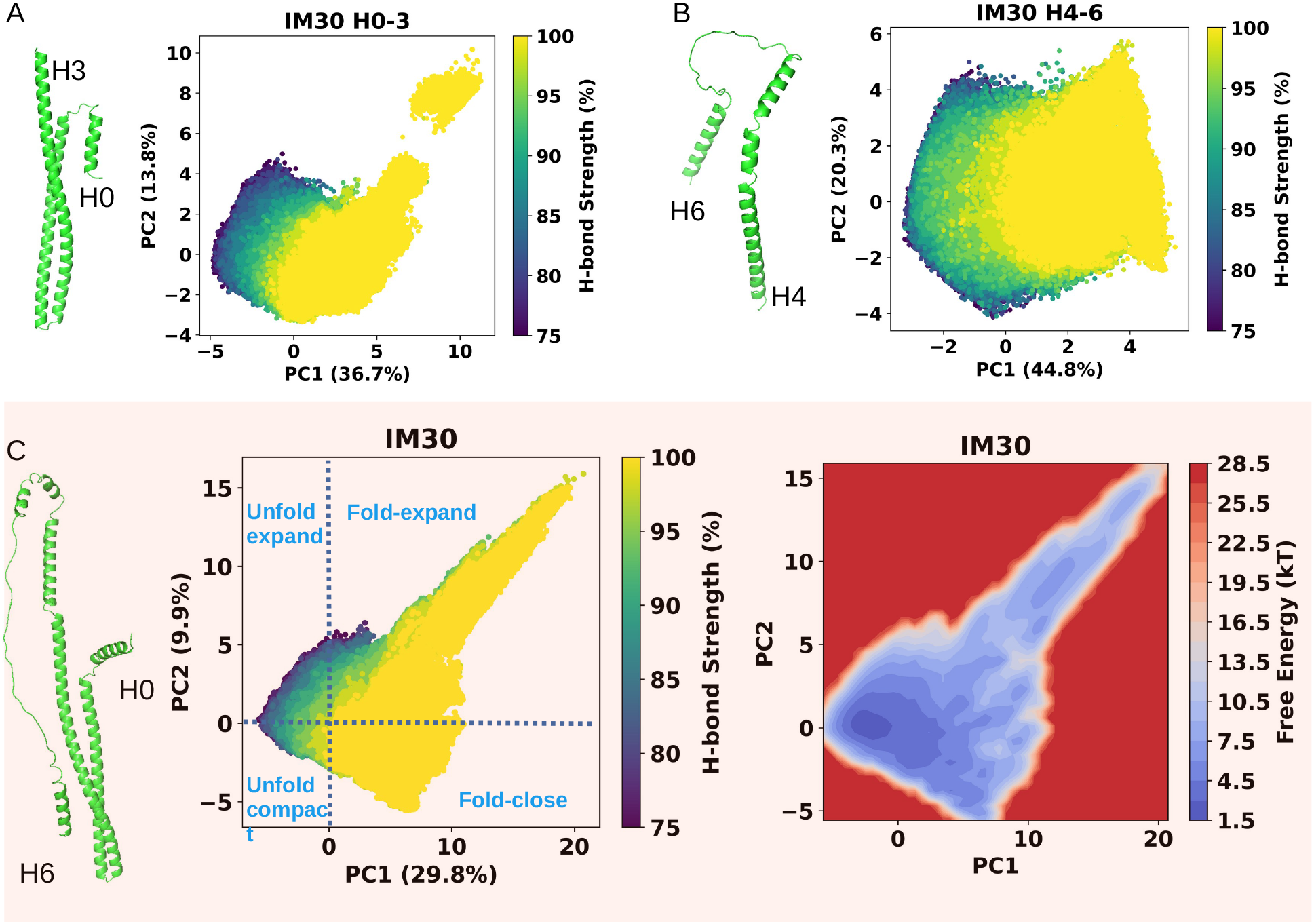
(A) PCA of IM30 H0-3 conformational ensembles, colored according to HBS. (B-C) PCA of IM30 H4-6 (B) and full-length IM30 (C), with conformations colored according to HBS. The left panels of all systems show representative conformations predicted by AlphaFold. (C) additionally presents the free energy landscape derived from the probability distribution of the conformational data for full-length IM30.

At lower HBS values, the conformational ensembles of H0-3 and H4-6 occupy similar regions of PCA space, reflecting comparable levels of compactness and disorder. However, with increasing HBS, H0-3 exhibits distinct population shifts and additional clusters associated with deviations from the dominant compact hairpin conformation. Correspondingly, both fragments display similar asphericity values below 95% HBS (Figure 3D), indicating comparable overall compactness under weaker hydrogen-bonding conditions. At higher HBS, H0-3 exhibits increased asphericity due to the formation of more extended helical hairpin conformations. The isolated cluster observed in the highly helical and expanded region of H0-3 corresponds to a coiled-coil like state (Figure 3F), distinct from the compact and transiently helical conformations predominantly populated at lower HBS. The redistribution of populations in PCA space is consistent with the trends observed in the radius of gyration (*R*_*g*_), further supporting the coupling between helix stabilization and global conformational reorganization.

### IDRs Enable a Continuum of Hairpin Conformations

The free-energy landscapes derived from the PCA distributions further illustrate how transient helicity regulates the conformation of IM30 and its constituent domains (Figure 4C (for IM30) and S8A-B). At low HBS, all constructs predominantly occupy disordered or weakly structured conformations, producing dense PCA clustering and shallow free-energy minima. Increasing HBS progressively stabilizes transient helices, reshaping the energy landscape and shifting populations toward more ordered and expanded conformational states. The H4-6 region exhibits broad and shallow free-energy basins, reflecting its lower intrinsic helical propensity and highly flexible polymer-like behavior. These shallow minima generate a rugged conformational landscape that enables extensive exploration of structural states rather than confinement within a single dominant basin. In contrast, H0-3 displays deeper and more localized energy minima in both cluster associated with stabilization of the conserved hairpin motif. However, when incorporated into full-length IM30, the flexible H4-6 region modulates these energetic preferences, preventing complete restriction to a single rigid state and instead promoting a continuous distribution of intermediate conformations. This coupling between transient helix formation and flexible flanking regions provides a mechanism for structural rearrangement within IM30. Transient helices continuously form and dissolve, while the disordered H4-6 region introduces additional conformational variability that facilitates transitions between neighboring states. Consequently, the full-length protein samples a broader and more continuous conformational landscape compared with isolated domains, allowing the conserved hairpin motif to maintain structural integrity while adapting to changes in its surrounding environment.

To further examine this effect, we analyzed the PCA landscape of the H0-3 region extracted from the full-length IM30 trajectory while excluding contributions from H4-6 (Figure 5). Unlike isolated H0-3, which exhibits a distinct population cluster corresponding to stabilized hairpin conformations, H0-3 within the full-length architecture samples a continuous range of conformations. This observation demonstrates that flanking IDRs do not simply increase disorder; rather, they redefine the accessible conformational space and facilitate continuum transitions of conserved structural motifs. Together, PCA and free-energy analyses demonstrate that HBS-dependent modulation of helicity does not produce a simple binary disorder to order transition. Instead, it generates a continuous conformational spectrum involving folded-expanded, unfolded-expanded, unfolded-compact, and folded-close states (Figure 4 and Figure 5). The resulting structural plasticity emerges from the balance between transient local helix stabilization and the flexibility introduced by IDRs, enabling conserved domains within modular ESCRT-III proteins to dynamically reorganize while preserving their overall architecture.

**Figure 5:**
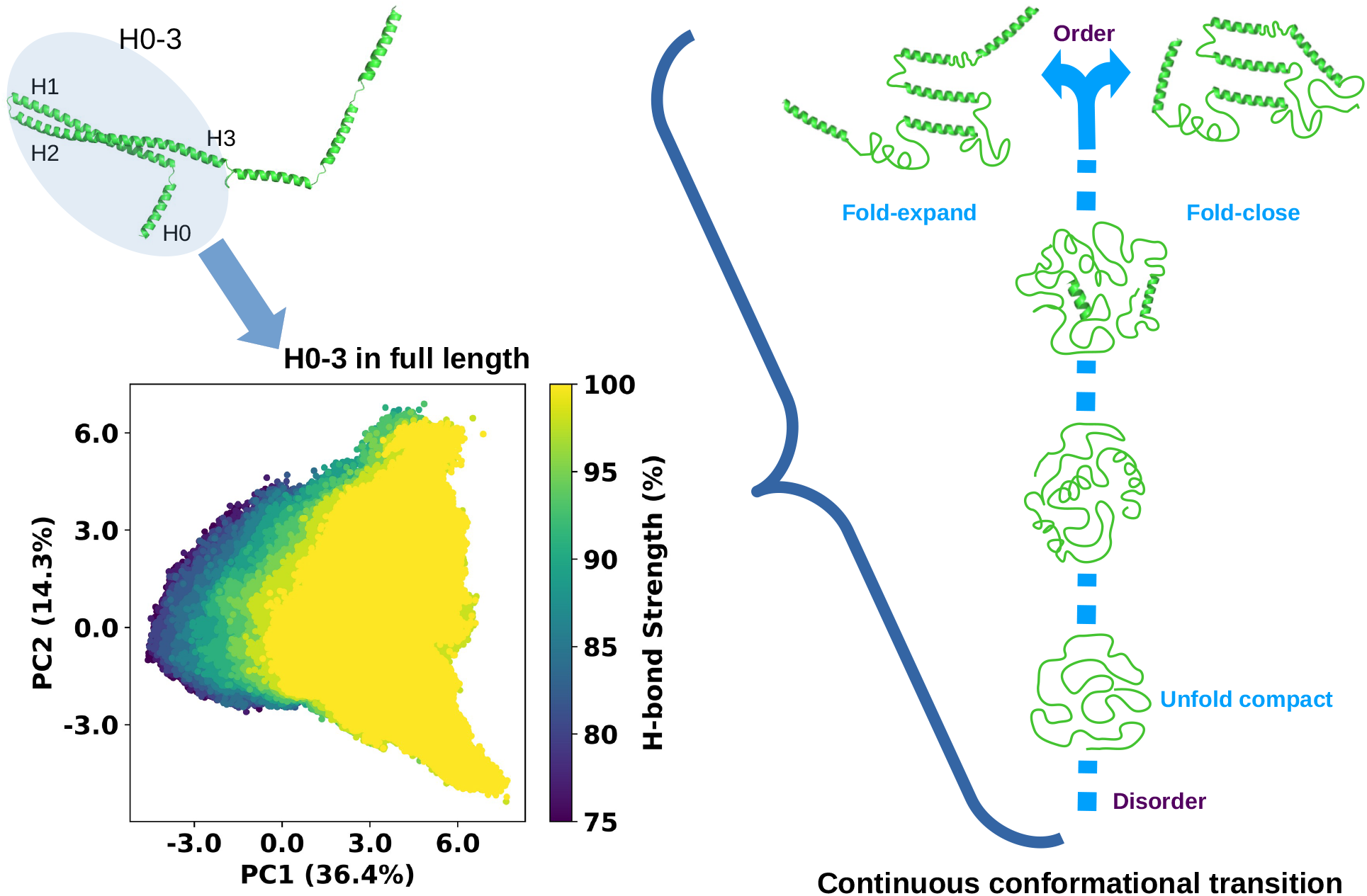
PCA of the H0-3 region within full-length IM30, colored according to HBS, showing a continuous conformational transition represented by the schematic diagram.

Collectively, helicity analysis, polymer-based structural characterization, PCA, and free-energy landscapes reveal that IM30 achieves a broad conformational ensemble through the interplay between structural order and disorder. The emergence of transient helices in the HBS crossover regime provides local structural stabilization, while flexible regions contribute to ensemble expansion by forming transient helical interactions. Together, these effects generate a continuous distribution of conformational states rather than discrete structural transitions, as illustrated in the right panel of Figure 5 for all modular protein system.

## Discussion

Structural flexibility is a fundamental determinant of the dynamic behavior and biological function of modular protein systems. Many proteins combine conserved structured elements with IDRs, allowing transitions between ordered, partially ordered, and disordered conformational states that regulate assembly, remodeling, and higher-order organization. However, the molecular principles by which disordered regions modulate the conformational behavior of conserved structural motifs remain challenging to characterize due to the heterogeneous and dynamic nature of their ensembles. In this work, we introduce a computational framework that systematically probes the continuum between ordered and disordered states by tuning HBS, an effective parameter controlling secondary-structure stabilization. Although HBS does not correspond to a single physiological variable, it provides a coarse-grained representation of changes in effective stabilizing interactions that may arise from environmental factors such as ionic conditions, molecular crowding, ligand binding, or membrane association. By varying HBS, we investigate how changes in local structural stability reshape the global conformational organization of IM30 and its modular components.

A major advantage of this framework is the combination of HREM with an efficient multi-chain CG simulation strategy. By simultaneously evolving multiple independent protein chains within a single dilute simulation box while preventing intermolecular interactions, this approach generates statistically robust conformational ensembles without requiring numerous independent simulations. The resulting large-scale datasets provide an efficient platform for ensemble-based structural analysis and future integration with data-driven approaches for investigating protein disorder, transient secondary structures, and sequence-structure relationships.

Beyond sampling efficiency, the HBS-based framework enables systematic characterization of transient helices, which represent important intermediate structural states that are often difficult to capture using conventional simulations. Continuous modulation of HBS allows exploration of ensembles spanning highly disordered conformations to locally stabilized helical states, revealing how transient secondary structures emerge and influence global organization. Rather than behaving as a binary switch between folded and unfolded states, proteins sample a continuous spectrum of conformations in which transient helices provide localized stabilization while maintaining ensemble diversity.

Application of this framework to the ESCRT-III protein IM30 demonstrates the biological relevance of transient helices in regulating modular protein architectures. IM30 contains a conserved helical hairpin scaffold flanked by intrinsically disordered regions, providing an ideal system for examining the interplay between structural order and disorder. Our results show that H0-3 possesses higher intrinsic helical propensity and stronger local ordering, whereas the H4-6 region exhibits broader conformational sampling characteristic of IDRs. Importantly, the disordered regions do not simply destabilize the conserved hairpin; instead, IDR-associated transient helices reshape its conformational landscape by reorganizing the states while maintaining the integrity of the core architecture. Compared with the isolated hairpin, the hair-pin within full-length protein explores more continuous conformational ensemble, demonstrating how flexible regions associated with conserved motifs regulate structural diversity.

Together, these findings establish a physical framework in which modulation of effective hydrogen-bond interactions controls the emergence of transient secondary structures and regulates the conformational landscapes of modular proteins. The results highlight that structural adaptation does not arise from a simple disorder to order transition, but from a dynamic balance between transient ordering and conformational diversity. More broadly, the combination of scalable ensemble sampling, physics-based perturbation, and multilevel structural analysis provides a general strategy for understanding how local interaction strengths and intrinsically disordered regions regulate protein conformational organization and functional versatility.

## Methods

### Coarse-Grained Model

All simulations were performed using a hybrid coarse-grained representation based on the PLUM force field [62]. This model preserves backbone geometry and captures secondary-structure formation while employing an implicit solvent. Each amino acid, except glycine, is represented by four beads: N, C*α*, C’, and C*β*, corresponding respectively to the amide nitrogen, central carbon, carbonyl carbon, and side chain. Glycine lacks the C*β* bead, and proline is treated using modified backbone parameters to account for its ring constraints. A key advantage of this model is the presence of an explicit backbone hydrogen-bond potential, which enables realistic helix formation even within an implicit-solvent description. The hydrogen-bond potential is given by

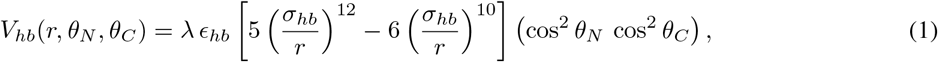

where *r* is the distance and *θ*_*N*_ and *θ*_*C*_ are the corresponding angular terms. The interaction is nonzero only when |*θ*_*N*_|, | *θ*_*C*_| *<* 90°. The bias parameter *λ ≥* 1 was varied across 12 replicas spanning the range 0.75 *≤ λ ≤* 1.0, allowing the system to undergo folding-unfolding transitions by effectively scaling the strength of secondary-structure formation.

### Simulation Framework

Initial configurations were generated using HOOBAS [94], and all systems were started from fully extended and structurally disordered conformations. Simulations were carried out using HOOMD-blue [95, 96] in a cubic box of volume 32000 Å^3^ to maintain dilute conditions among the 64 chains. An initial relaxation run of 100 ns was performed in the microcanonical (NVE) ensemble with a displacement cap of 0.02. Production runs were then carried out in the canonical (NVT) ensemble using Langevin dynamics with a timestep of *δt* = 0.01 *τ*, corresponding to 10 fs. To enhance sampling efficiency, the viscosity (friction) coefficient was reduced from 1 to 0.001 compared to our previous studies [17]. This modification reduces the friction timescale, thereby accelerating dynamical relaxation while preserving the correct equilibrium distribution ensured by Langevin dynamics. Enhanced sampling was performed using HREM along the *λ* space [67, 97, 98]. Chains were shuffled among replicas prior to each exchange attempt to enhance sampling efficiency and promote thorough exploration of phase space. Data handling and workflow management for the simulations were carried out using signac and signac-flow [99–101]. Visualization were performed with OVITO [102].

Autocorrelation functions computed for *R*_*g*_ and *R*_*ee*_ yield a correlation time of approximately 2 *µ*s, with continuous decay. To ensure at least ten statistically independent samples, each trajectory was therefore simulated for a total of 20 *µ*s. Following inspection of the temporal evolution of structural observables (e.g., *R*_*g*_ and *R*_*ee*_, *A*3), the first 10 *µ*s were discarded to eliminate transient effects associated with equilibration. This left 10 000 equilibrated frames for analysis in each trajectory. The full-length IM30 protein, its N-terminal with coiled-coil region (H0-3), and its C-terminal region (H4-6) were simulated independently using the same HREM protocol and simulation parameters.

## Additional Information

Further details on system equilibration, HREM, structural property analysis, and PCA are provided in the Supporting Information.

## Acknowledgments

We thank Prof. Friederike Schmid (Institute of Physics, Johannes Gutenberg University Mainz) for carefully reading the manuscript and for her valuable comments and suggestions.

## Author Contributions

T.R.B. and M.G. conceived the study and developed the methodology. T.R.B. performed the simulations and carried out the data analysis. T.R.B. and M.G. wrote the original draft of the manuscript. M.G. and K.K. contributed to reviewing and editing the manuscript. M.G. acquired funding and administered the project. M.G. and K.K. supervised the research. K.K. provided computational resources and infrastructure.

## Data Availability

All code and data used in this study will be made available upon publication of the final manuscript.

