## Supporting Information for "Multi-Chain Hamiltonian Replica Exchange Framework for Probing Transient Helices and Order-Disorder Transitions in Proteins"

### Supplementary Information

#### Hamiltonian Replica Exchange Method (HREM)

To enhance sampling across conformational space, we employed a two-dimensional Hamiltonian replica exchange scheme [1–3]. The first exchange dimension corresponded to the bias parameter $\lambda$, which modulates the strength of the backbone hydrogen-bond potential; the second dimension involved reshuffling the identities of the 64 simulated chains across replicas at fixed $\lambda$ values. The 64 chains were positioned on a $4 \times 4 \times 4$ lattice of anchor points to avoid steric overlap between chains. Replica exchange attempts were performed every $10^5$ integration steps. Prior to each attempt, chains were randomly reassigned among replicas. Following an accepted Metropolis exchange of force-field parameters, the corresponding chain was translated equal to displacement to avoid unintended spatial overlaps arising from the reassignment procedure and to preserve excluded-volume interactions. Exchange attempts were evaluated using the Metropolis acceptance probability [4],

$$P_{\text{acc}} = \begin{cases} 1, & \Delta E < 0, \\ \exp(-\Delta E), & \Delta E \geq 0, \end{cases} \tag{1}$$

implemented numerically as

$$\text{accept if} \quad \text{random} < \min\big(1, e^{-\Delta E}\big) \, .$$

This ensures detailed balance and correct equilibrium sampling across the bias ladder. The observed exchange success rates ranged from 10% to 80%, indicating efficient replica diffusion and proper functioning of the HREM scheme.

#### Replica Exchange Equilibration

The sequences of IM30, H0-3 and H4-6 (given in S8 was used to generate the initial configuration for simulation. Snapshots of IM30 are shown in Figure S2A. These conformations are extracted from the simulation with 100% HBS. The extended structure is mostly observed at non-equilibrium, while overlapped globular is seen frequently after equilibrium. For the equilibrated system, the constant H-bond potential for a given bias ((Figure S1A) and monotonic behaviors of the overall potential varying with bias (Figure S1B), and the distribution of exchange acceptance rates (Figure S1C-D) indicate that the HREM was successfully implemented in the simulation. Longer chain and higher biasing strength have a greater (more negative) H-bond potential as compared to other system. For the same conditions higher success rate of exchange was recorded for H0-3 and H4-6 as compared to IM30.

IM30 is a structured protein which takes a long time for equilibration, thus this was monitored by plotting the evolution of structural features such as radius of gyration ($R_g$), end to end distance ($R_{ee}$), as well as asphericity ($A3$). The $R_g$ provides a measure of the polymer's overall mass distribution around its center of mass, reflecting how compact or spread out the chain is. The $R_{ee}$ measures the linear distance between the two ends of the polymer, indicating the degree of chain extension, and $A3$ measures the degree of spherical symmetry (a perfect sphere has $A3$ equal to zero) [5, 6]. Together, these metrics provide complementary insights into the polymer's conformational state and dynamics. Starting from fully disordered and extended conformations, the structural evolution of the chains over time is shown in Figures S2C-E and S3A-F, for all biases. As the strength of the hydrogen bond decreases, the curve exhibits a faster decay and a steeper slope. These decay curves for $R_g$ and $R_{ee}$ were further fitted exponentially to get the correlation time ($\tau$). When the chains are more structured and stiff, the above properties of polymeric chains tend to be more highly correlated, reflecting reduced flexibility and more extended conformations. The coefficients for the correlation from evolution of $R_g$ and $R_{ee}$ are reported in Figure S2 B. IM30 at 100% HBS has comparable highest values of $\tau$ as 1.89 $\mu$s and 1.76 $\mu$s for $\tau_{R_g}$ and $\tau_{R_{ee}}$ respectively. Both the values demand roughly 20 $\mu$s for full length simulation to capture at least ten uncorrelated data points. Also, evolution of $R_g$, $R_{ee}$ and $A3$ shows that, chain is equilibrated at $\approx 10 \mu$s. Therefore, data from the equilibration period were discarded, and only post equilibration data were used for analysis.

### Structural properties

The structural properties of the chains were analyzed by calculating the radius of gyration ($R_g$), the asphericity from the gyration tensor ($A_3$), the end-to-end distance ($R_{ee}$), the persistence length ($l_p$), and the autocorrelation function ($C(t)$) [5, 6]. The radius of gyration provides a measure of the overall size of the polymer and is defined as

$$R_g^2 = \frac{1}{N} \sum_{i=1}^{N} |\mathbf{r}_i - \mathbf{r}_{\mathrm{cm}}|^2, \tag{2}$$

where $N$ is the number of monomers, $\mathbf{r}_i$ is the position vector of the $i$-th monomer, and $\mathbf{r}_{\mathrm{cm}}$ is the center of mass of the chain. To quantify the overall shape anisotropy of each chain, we computed the asphericity parameter $A_3$ from the gyration tensor. The gyration tensor $Q$ was then calculated from the position vectors relative to the chain's center of mass. Diagonalization of $Q$ yields its eigenvalues $\lambda_1, \lambda_2, \lambda_3$, which characterize the principal spatial extensions of the chain. The asphericity was computed as

$$A_3 = \frac{1}{6} \frac{\sum_{i=1}^{3} (\lambda_i - \bar{\lambda})^2}{\bar{\lambda}^2}, \qquad \bar{\lambda} = \frac{\lambda_1 + \lambda_2 + \lambda_3}{3}, \tag{3}$$

which measures the deviation of the chain from spherical symmetry ($A_3 = 0$ for a perfectly spherical distribution). The $R_{ee}$, which measures the distance between the first and last monomers of the chain, is defined as

$$R_{ee} = |\mathbf{r}_N - \mathbf{r}_1|. \tag{4}$$

The persistence length, which characterizes chain stiffness, is obtained from the decay of angular correlations along the backbone as

$$\langle \cos\theta(s) \rangle = \exp(-s/l_p), \tag{5}$$

where $\theta(s)$ is the angle between bond vectors separated by contour distance $s$. Finally, the autocorrelation function, describing the time-dependent correlation of a property $X$ (such as $R_g$ or $R_{ee}$), is defined as

$$C(t) = \frac{\langle X(0)X(t) \rangle - \langle X \rangle^2}{\langle X^2 \rangle - \langle X \rangle^2}. \tag{6}$$

The decay of $C(t)$ provides an estimate of the relaxation time of the chain.

### Cluster Analysis via PCA

For the cluster analysis, we first calculated the helical propensity of each helical segment of the respective chains over time, considering seven segments for the full-length protein, four segments for the coiled-coil region, and three segments for the C-terminal region. In addition, we computed the pairwise distances between the centers of mass of these segments. These quantities were combined into feature vectors for each sampled conformation:

$$\mathbf{x}^{(k,b)} = \left[ p_1^{(k,b)}, p_2^{(k,b)}, \ldots, p_M^{(k,b)}, d_{12}^{(k,b)}, d_{13}^{(k,b)}, \ldots, d_{MN}^{(k,b)} \right]^T, \tag{7}$$

where $p_i^{(k,b)}$ is the helical propensity of segment $i$ in conformation $k$ under bias $b$, $d_{ij}^{(k,b)}$ is the distance between segments $i$ and $j$, $M$ is the total number of helical segments, and $N$ is the number of segment pairs.

To ensure all features contributed equally, the feature vectors were standardized using Z-score normalization:

$$\tilde{x}_i^{(k,b)} = \frac{x_i^{(k,b)} - \mu_i}{\sigma_i}, \tag{8}$$

where $\mu_i$ and $\sigma_i$ are the mean and standard deviation of feature $i$ across all conformations and biases.

Principal component analysis (PCA) [7] was then performed on the standardized data. Let $\mathbf{X} \in \mathbb{R}^{K\times F\times B}$ be the 3-dimensional matrix of standardized feature vectors ($K$ conformations per bias, $F$ features, $B$ bias values). The principal components are obtained by flattening the bias dimension when analyzing all biases together:

$$\mathbf{Y} = \mathbf{X}_{\text{flat}}\mathbf{W}, \tag{9}$$

where $\mathbf{X}_{\text{flat}} \in \mathbb{R}^{(K\cdot B)\times F}$ is the reshaped matrix, $\mathbf{W} \in \mathbb{R}^{F\times L}$ contains the eigenvectors of the covariance matrix

$$\mathbf{C} = \frac{1}{K\cdot B - 1}\mathbf{X}_{\text{flat}}^T\mathbf{X}_{\text{flat}}, \tag{10}$$

and $L$ is the number of principal components retained. The first eigenvalue $\lambda_1$ represents the variance captured by the dominant mode. The resulting PCA scores $\mathbf{Y}$ were binned and used to generate heatmaps of the average replica values, highlighting the distribution of conformations across PCA space. The contribution of each feature to the principal components was quantified via the PCA loadings:

$$\mathbf{L} = \mathbf{W}\cdot\sqrt{\mathbf{\Lambda}}, \tag{11}$$

where $\mathbf{\Lambda}$ is the diagonal matrix of eigenvalues. PCA was performed both on the combined dataset including all bias values, allowing identification of clusters of conformations with similar structural characteristics and their relation to the underlying helical and spatial organization of the protein segments.

### Supporting Figures

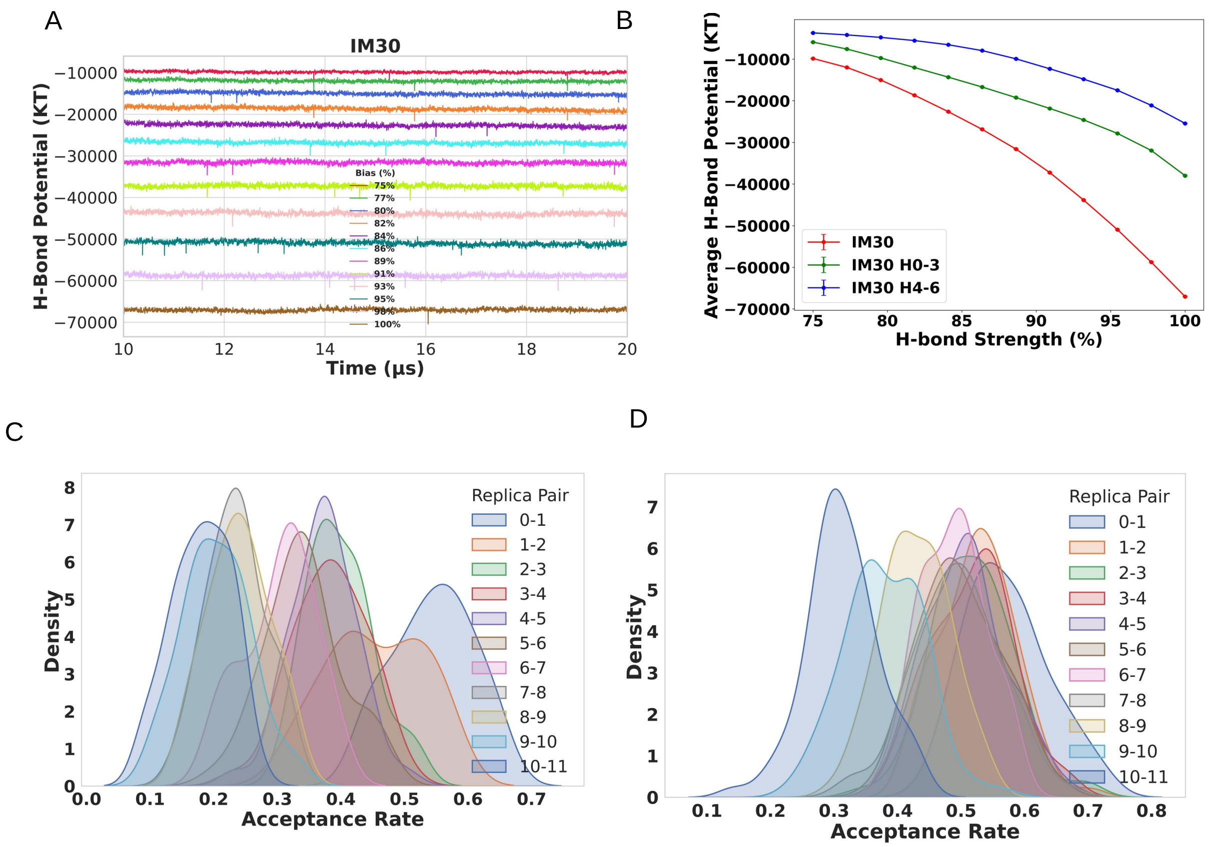


Figure S1: (A) Time-resolved hydrogen-bond potentials for IM30, H0-3, and H4-6 across varying H-bond bias strengths.(B) Hydrogen-bond potentials for the four systems.(C–D) Probability distributions of exchange acceptance ratios between adjacent Hamiltonian replicas for IM30 (C) and H0-3 (D).

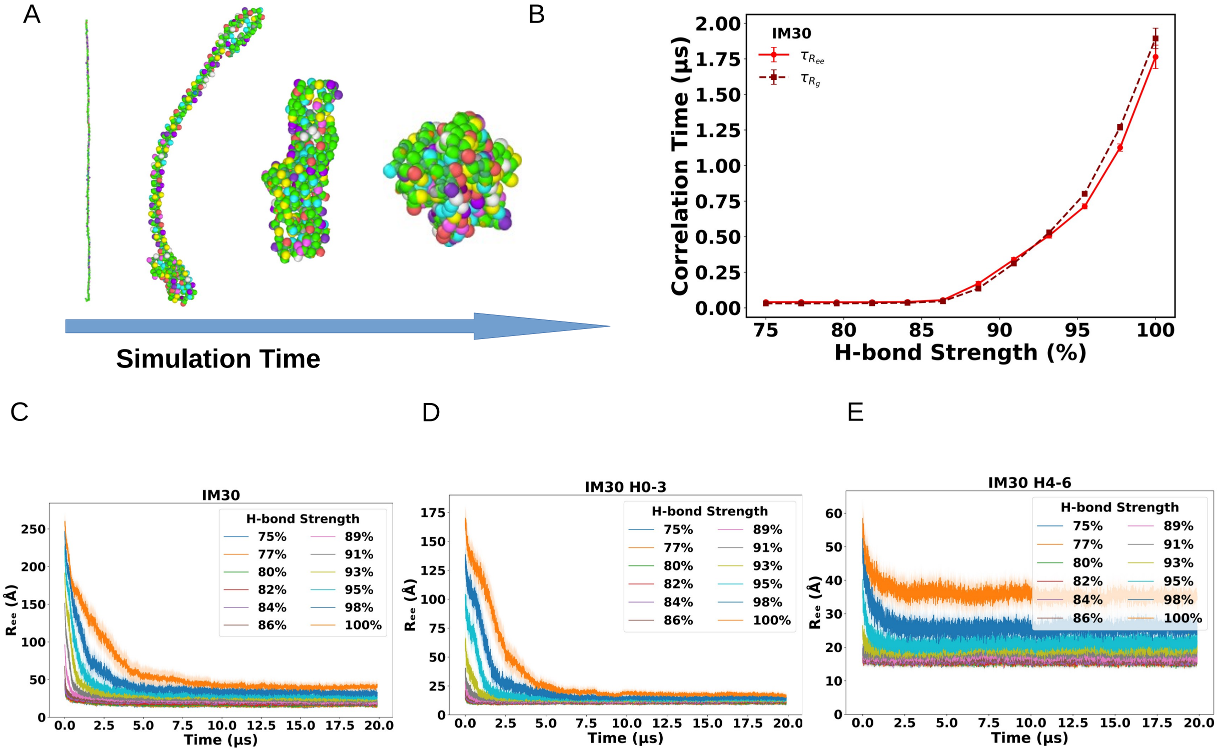


Figure S2: (A) Snapshots of IM30 at different time frame of simulation. (B) Variation of correlation time with H-bond strength calculated from the exponential fitting of time series data of $R_g$ and $R_{ee}$. (C-E) End-to-end distance ($R_{ee}$) as a function of time for IM30 (C), H0-3 (D), and H4-6 (E) under the same bias conditions, likewise exhibiting exponential decay.

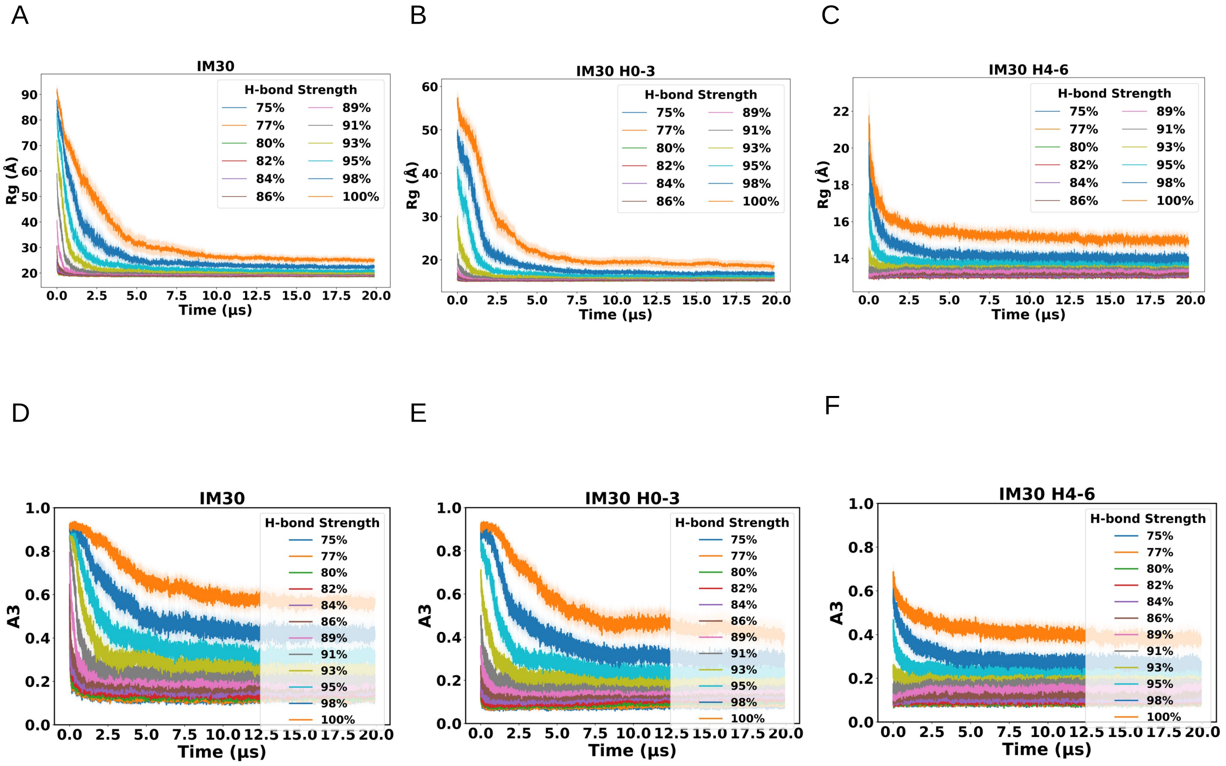


Figure S3: (A-C) Radius of gyration ($R_g$) as a function of time for all bias conditions for IM30 (A), H0-3 (B), and H4-6 (C), highlighting an exponential decrease in compactness.(D–F) Asphericity ($A3$) as a function of time for IM30 (D), H0-3 (E), and H4-6 (F) under the same bias conditions representing deviation of chains from extending to spherical.

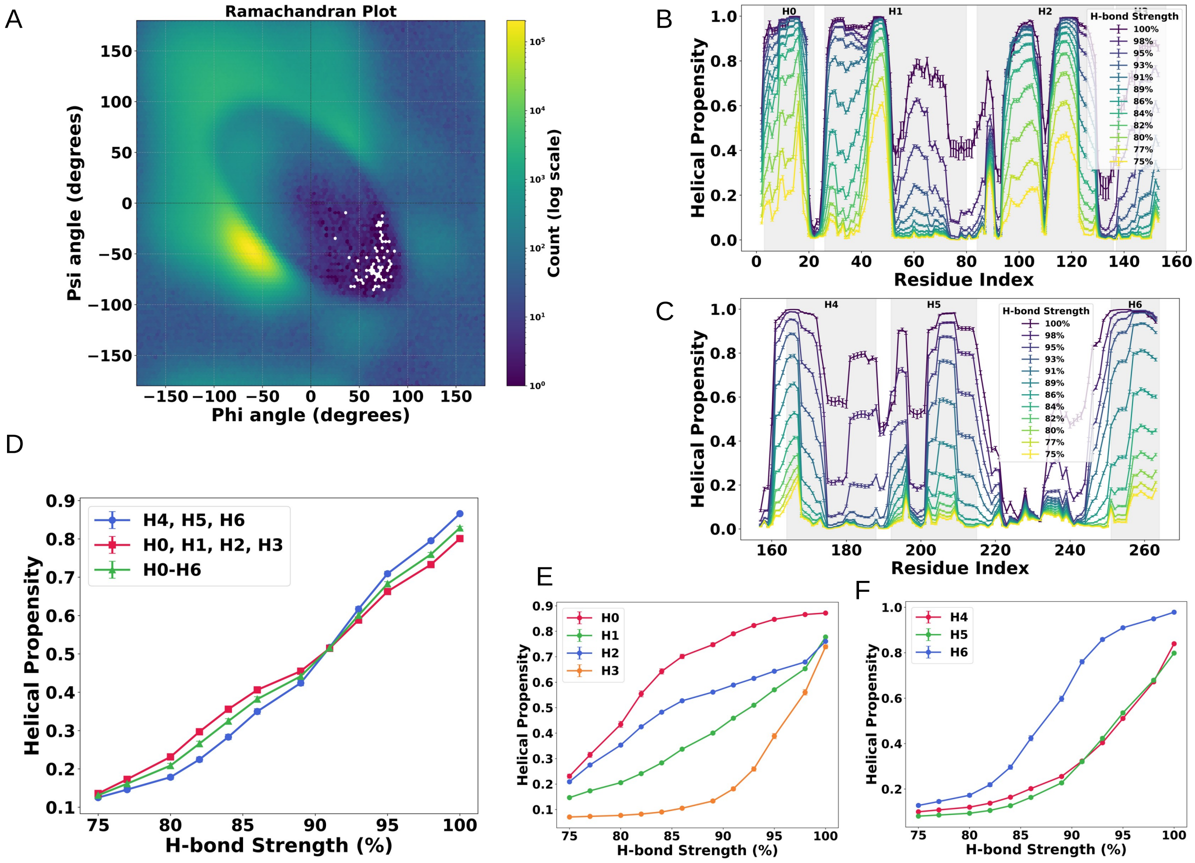


Figure S4: (A) Ramachandran plot illustrating the $\alpha$-helical region defined by clustered backbone dihedral angles of IM30. (B-C) Helical propensity profiles for H0-3, and H4-6 across different H-bond bias strengths (100 % shown in dark, 75 % shown in yellow). (D)Average helicity of H0,H1,H2,H3 segment, H4,H5,H6 segments of IM30 (E) Average helical propensity of the four N-terminal helices (H0-H3) in H0-3. (F) Average helical propensity of the four C-terminal helices (H4-H6) in H4-6, computed from panels B and C.

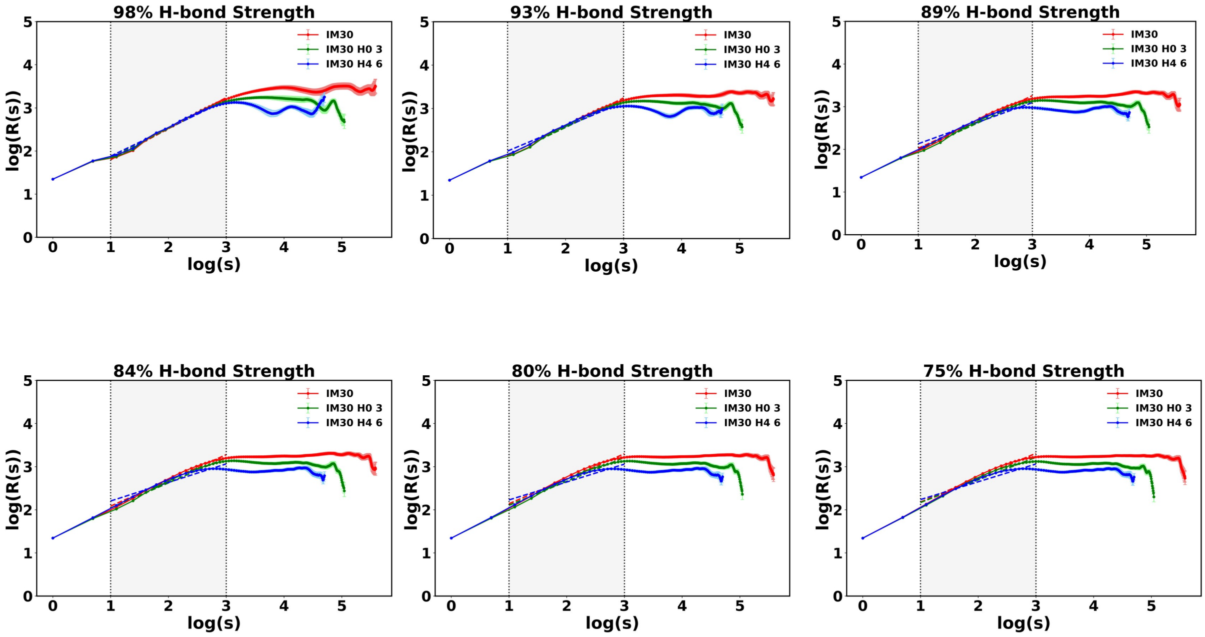


Figure S5: Logarithmic-scale plots of R(s) with residue spacing for IM30, H0-3, and H4-6 across the full range of H-bond bias strengths. Data for IM30, H0-3, and H4-6 are shown in red, green, and blue, respectively, with dotted lines marking the fitted scaling regimes.

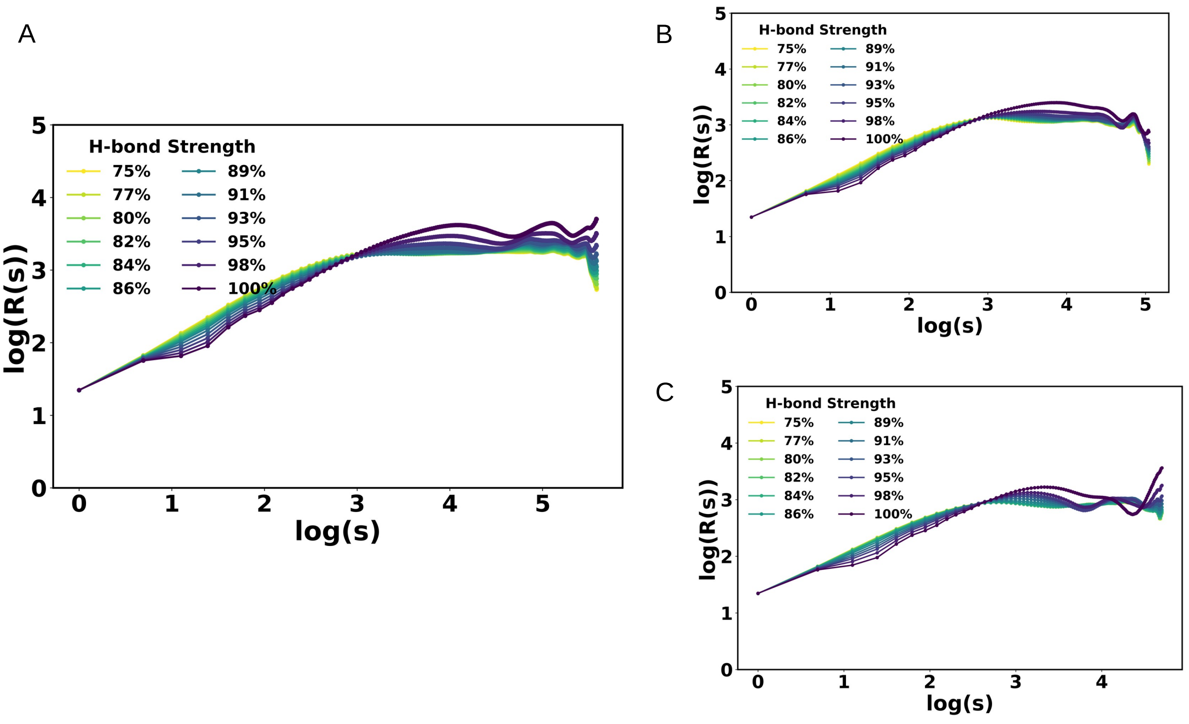


Figure S6: Log-log scaling relations of $R(s)$ with residue spacing for (A) IM30, (B) H0-3, and (C) H4-6 under different H-bond bias strengths. Data obtained at 100 % bias are shown in dark tones, whereas those at 75 % bias are shown in yellow.

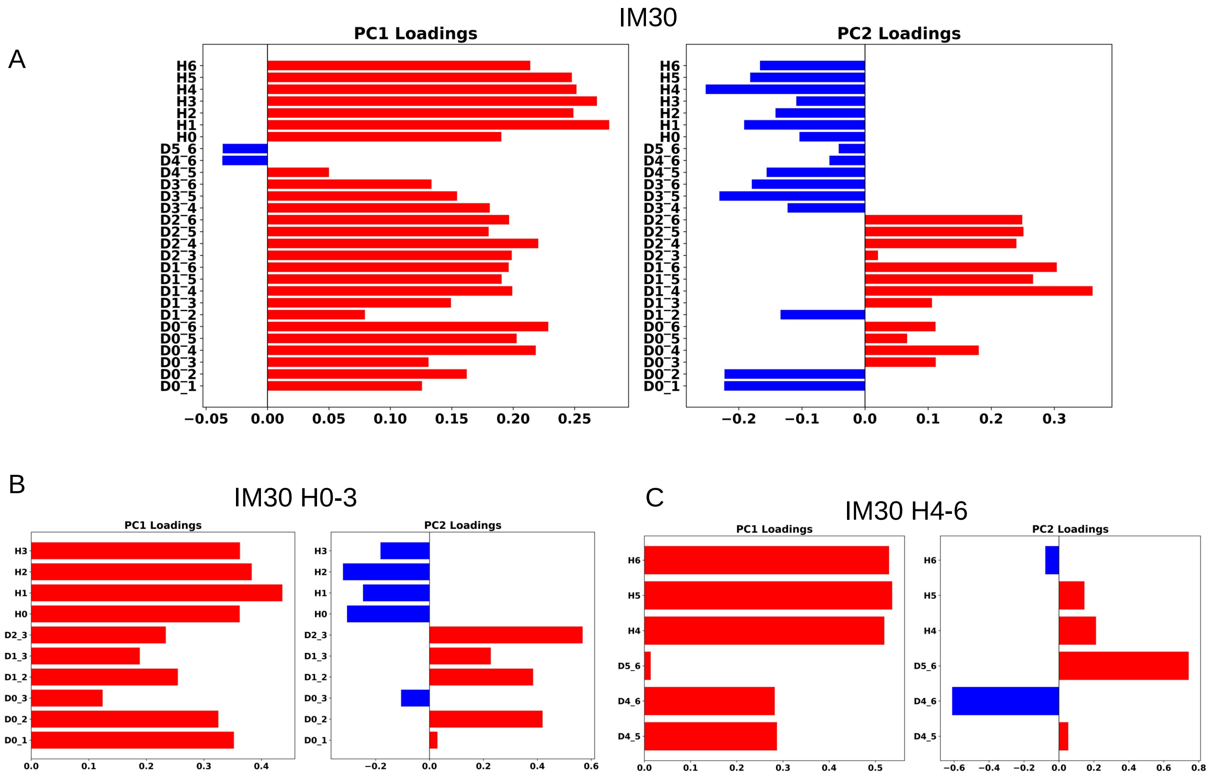


Figure S7: (A-C)Contribution of features for PCA of the system as loadings for IM30 (A), H0-3 (B), and H4-6 (C) respectively.

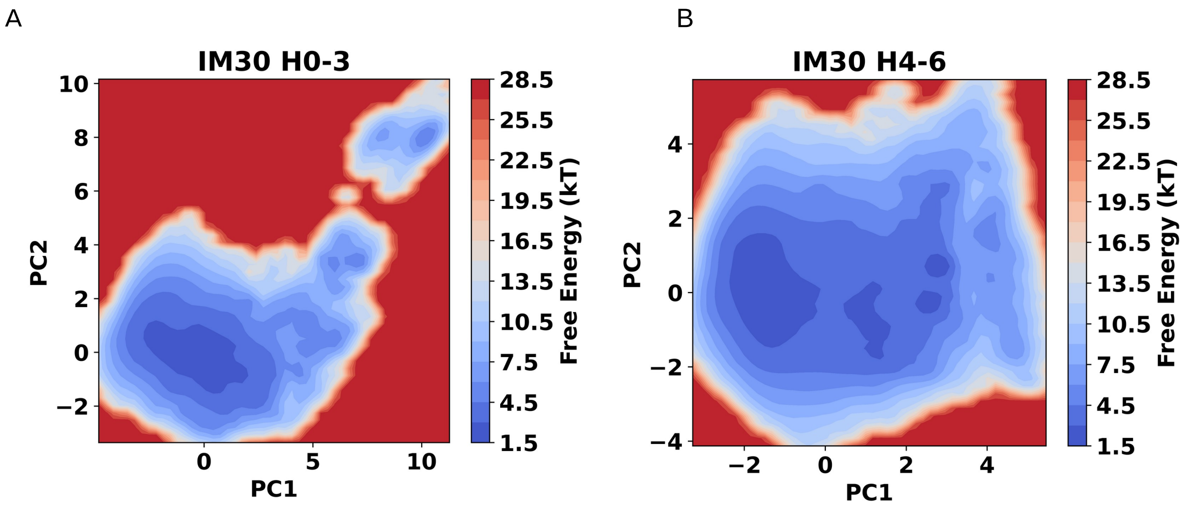


Figure S8: (A) Principal component analysis (PCA) of PCA of H0-3 (B) and H4-6, colored according to free energy computed from the probability distribution (kT=1) highlighting differences in conformational distribution and stability.

**IM30:**

MGLFDRLGRVVRANLNDLVSKAEDPEKVLE**QAVIDMQEDLVQLRQAVARTIAEEKRTEQRLNQDTQEA KKWEDRAKLALT**NGEENLAREALARKKSLTDTAAAYQTQLAQQRTMSENLRRNLAALEAKISEAKTKK NMLQARAKAAKANAELQQTLGGLGTSSATSAFERMENKVLDMEATSQAAGELAGFGIENQFAQLEAS SGVEDELAALKASMAGGALPGTSAATPQLEAAPVDSSVPANNASQDDAVIDQELDDLRRRLNNL

**IM30 H0-3:**

MGLFDRLGRVVRANLNDLVSKAEDPEKVLEQAVIDMQEDLVQLRQAVARTIAEEKRTEQRLNQDTQEA KKWEDRAKLALTNGEENLAREALARKKSLTDTAAAYQTQLAQQRTMSENLRRNLAALEAKISEAKTKKN MLQARAKAAKANAELQQTL

**IM30 H4-6:**

GGLGTSSATSAFERMENKVLDMEATSQAAGELAGFGIENQFAQLEASSGVEDELAALKASMAGGALP GTSAATPQLEAAPVDSSVPANNASQDDAVIDQELDDLRRRLNNL

Figure S9: Aminoacid sequences for IM30 (Highlight for H1), H0-3 and H4-6 of IM30

[4] Daan Frenkel and Berend Smit. *Understanding molecular simulation: from algorithms to applications*. Elsevier, 2023.

[5] Michael Rubinstein and Ralph H Colby. *Polymer physics*. Oxford university press, 2003.

[6] Viktoria Blavatska and Wolfhard Janke. Shape anisotropy of polymers in disordered environment. *The Journal of chemical physics*, 133(18), 2010.

[7] Ian Jolliffe. Principal component analysis. In *International encyclopedia of statistical science*, pages 1094–1096. Springer, 2011.
